# Biomes do not show a clear-cut phylogeny

**DOI:** 10.1101/2025.07.02.662740

**Authors:** Gaucherel Cédric, Noûs Camille, Hély Christelle

**Author notes:** <u>Correspondence to:</u> Cédric Gaucherel, INRAE – ECODIV, UMR AMAP, TA A.51/PS2, 34398 Montpellier Cedex 5 (France).

## Abstract

Shifts between biomes, the broad types of ecosystems, are far less understood than other ecological transitions. In this study, we aimed at identifying a possible and preliminary lineage between terrestrial biomes on Earth by using up-to-date phylogenetic methods. Although a deep statistical power was lacking, we built on expert knowledge a double-entry table filled by a variety of traits characterizing all the 14 terrestrial biomes on Earth. As a central result, the biome phylogeny computed here is a clear approximation of the dominant biome successions observed on Earth. Latitudes (i.e., locations), not included into the trait table, may first explain the computed phylogeny. In particular, tropical biomes appeared well related and exhibited a possible common ancestor (higher than 50% confidence level). We discussed this possible history of terrestrial biome on Earth, along to the involved processes over the long term. This helped explaining how biomes differ from purely biological materials and partly why ecosystems are not evolving as life does.

## Introduction

Our planet is covered by a diversity of broad regional conditions, mainly in terms of climate, flora and fauna. Such regions are called biomes and are represented by broad ecosystem types. Biomes have historically shifted, due to a variety of processes such as plate tectonics, climatic conditions and, more recently, human impacts [1-3]. Although we start locating and understanding ecosystem transitions, i.e., a possible shift between two distinct ecosystems [4, 5], shifts between biomes are far less understood [6-9]. Moreover, we lack a clear and integrated view of biome transitions along time, possibly leading to biome trajectories over the long term. In this study, we aimed at identifying a possible and preliminary lineage of terrestrial biomes on Earth by using up-to-date phylogenetic methods.

A range of observations in paleoecology confirms that, in many different locations on Earth, biomes have shifted from one to another, sometimes irreversibly [e.g., 10], sometimes back and forth [e.g., 11, 12-14]. Simultaneously, various models successfully reconstruct the past variations of biomes in most locations on Earth, and at various periods [e.g., 14, 15]. Conventionally, scientists identify 14 terrestrial and 25 contrasted biomes including aquatic ones, and start now understanding the processes and interactions responsible for every isolated biome transition [8, 16]. Yet, each biome transition involves two different biomes, although a broad integrated history would require grasping both successions of biomes (in time) and their neighborhood (in space). Few models are today equipped to grasp such biome transitions [17-19], which may be related to successions, disturbance impacts, or even environmental changes. It would be useful, at least, to identify possible successions in terms of composition at the biome level.

In other words, we question here a broad and integrated view of biome transitions over the long term, whatever the location on Earth. This question may be summarized into the need to identify biome lineages or, further, what may be called a “biome phylogeny”, if any. Many tools have been developed for building a phylogeny, as soon as the user assumes that units (biomes) are temporally inheriting from ancestors. Here, conversely to usual studies in phylogeny [6, 20], we did not assume that biomes have a unique lineage, as they do not possess a kind of DNA molecule gathering their constitutive information. Rather, we intended to identify one possible history of biome transitions among a range of possible histories. In addition, we did not perform such an analysis based on the species composition of each biome, but rather on generic traits identified by expert knowledge. This generic approach is often selected when phylogenetic tools are applied no to species lineage with purely biological units, but to other types of entities such as myths, rupestrian paintings or galaxies [21, 22]. Here, we only assumed that several biome transitions are possible, and we tried to list their proximity in terms of trait (character) compositions. Additionally, we did not possess a calibrated clock for biome transitions, although we assumed a regular one to broadly quantify similarities between biomes.

Hence, although a deep statistical power is lacking, we built a double-entry table filled by a variety of traits (properties) characterizing all the 14 terrestrial biomes on Earth, more based on expert knowledge than on data with a deep statistical power. We hypothesized (H1) that latitudes (i.e., locations, not included into the trait table) may first explain the computed phylogeny. In addition, we hypothesized (H2) that some filiations at specific latitudes (tropical versus temperate or higher latitudes) may be more robust and could explain well-known biome transitions. For example, tundra and boreal forests should be highly similar and may shift from one to the other. We then applied common phylogenetic tools on the table to identify the most probable (consensual) phylogeny of biomes and interpret identified filiations [23, 24]. We then discussed the possible history of terrestrial biome transitions on Earth and involved processes over the long term [8, 25], before discussing the assumptions, uncertainties and significance of this tentative phylogeny.

## Materials and methods

We used the 14 major terrestrial biomes conventionally defined by the WWF - Terrestrial Ecoregions of the World [Table 1, 16]. We added two biomes to root the phylogenetic tree: *Lake* and *RockIce* (representative of terrestrial lakes and non-vegetated (rocks and ice) areas, respectively) for which we computed the same variables than for the other biomes.

**Table 1.** List of the 14 major biomes from WWF - Terrestrial Ecoregions of the World [16], their cover (percentage of the total terrestrial surface) and abbreviation used in Fig. 1. Terrestrial lakes and bare soils have been retained for the rooting the phylogeny.

|  | Biome | Area (in %) | Abbreviation |
| --- | --- | --- | --- |
| 1 | Tropical and Subtropical Moist Broadleaf Forests | 13.4 | TrMBF |
| 2 | Tropical and Subtropical Dry Broadleaf Forests | 2.0 | TrDBF |
| 3 | Tropical and Subtropical Coniferous Forests | 0.5 | TrCoF |
| 4 | Temperate Broadleaf and Mixed Forests | 8.7 | TeBMF |
| 5 | Temperate Coniferous forests | 2.8 | TeCoF |
| 6 | Boreal Forests / Taiga | 10.3 | BoF |
| 7 | Tropical and subtropical grasslands, savannas, and shrublands | 13.7 | TrGSS |
| 8 | Temperate grasslands, savannas, and shrublands | 6.9 | TeGSS |
| 9 | Flooded grasslands and savannas | 0.7 | FIGS |
| 10 | Montane grasslands and shrublands | 3.5 | MoGS |
| 11 | Tundra | 7.9 | Tun |
| 12 | Mediterranean forests, woodlands, and scrub | 2.2 | MeFWS |
| 13 | Deserts and xeric shrublands | 18.9 | DeXS |
| 14 | Mangroves | 0.2 | Man |
|  | Continental lakes | 0.7 | Lake |
|  | Bare soils (rock and ice covers) | 7.5 | RockIce |

To characterize biomes, we gathered a total of 43 spatial indices (Table 2), from which we computed the median, the first and the ninth deciles per each biome, according to their extent (imprint) on Earth (Fig. 1). Bioclimatic variables are derived from the monthly temperature and rainfall values in order to generate more biologically meaningful variables [26]. The bioclimatic variables represent annual trends (e.g., mean annual temperature, annual precipitation) seasonality (e.g., annual range in temperature and precipitation) and extreme or limiting environmental factors (e.g., temperature of the coldest and warmest month, and precipitation of the wet and dry quarters, i.e., periods of three months). Soil properties are derived from the IGBP-DIS database at spatial resolution of 0.25° [27], and Gridded Population of the World, Version 4, with demography of year 2000 [28]. Elevation is derived from World digital elevation model EEA 1990, because SRTM data resampled was not available for polar latitudes [29]. We used biodiversity maps (i.e., the total species richness or total number of species present per location/pixel) and endemism maps (i.e., species having a unique and restricted spatial distribution smaller than 250,000 km^2^) for birds, amphibians and mammals only [8]. Finally, we used connectivity and Shannon indices for the Land/Ocean distribution (hypothesis of boundary influences), Shannon index for the continent fragmentation by different countries (hypothesis of country management influences), and Evenness index for land cover heterogeneity (hypothesis of modern land cover influences) computed on the basis of the MHM software [30-32].

**Table 2.** List of the variables used to characterize the 14 terrestrial biomes (see main text for associated references).

| Name | Description | Source |
| --- | --- | --- |
| **bio1** | Annual Mean Temperature | WorldClim |
| **bio2** | Mean Diurnal Range (Mean of monthly (max temp - min | WorldClim |
| **bio3** | Isothermality (BIO2/BIO7) (* 100) | WorldClim |
| **bio4** | Temperature Seasonality (standard deviation *100) | WorldClim |
| **bio5** | Max Temperature of Warmest Month | WorldClim |
| **bio6** | Min Temperature of Coldest Month | WorldClim |
| **bio7** | Temperature Annual Range (BIO5-BIO6) | WorldClim |
| **bio8** | Mean Temperature of Wettest Quarter | WorldClim |
| **bio9** | Mean Temperature of Driest Quarter | WorldClim |
| **bio10** | Mean Temperature of Warmest Quarter | WorldClim |
| **bio11** | Mean Temperature of Coldest Quarter | WorldClim |
| **bio12** | Annual Precipitation | WorldClim |
| **bio13** | Precipitation of Wettest Month | WorldClim |
| **bio14** | Precipitation of Driest Month | WorldClim |
| **bio15** | Precipitation Seasonality (Coefficient of Variation) | WorldClim |
| **bio16** | Precipitation of Wettest Quarter | WorldClim |
| **bio17** | Precipitation of Driest Quarter | WorldClim |
| **bio18** | Precipitation of Warmest Quarter | WorldClim |
| **bio19** | Precipitation of Coldest Quarter | WorldClim |
| **bulkdens** | bulk density (this variable only takes the integer values | IGIP-DIS 2000 |
| **Cac** | carbonate carbon density (in kg C/m<sup>3</sup> for 0-100cm depth | IGIP-DIS 2000 |
| **fieldcap** | fielcap (mm | IGIP-DIS 2000 |
| **nitrogen den** | total nitrogen density (g/m<sup>3</sup>) | IGIP-DIS 2000 |
| **pawc** | the available water capacity | IGIP-DIS 2000 |
| **ph1** | the soil pH between 0 and 30 cm depth | IGIP-DIS 2000 |
| **ph2** | the soil pH between 30 and 100 cm depth | IGIP-DIS 2000 |
| **soil carb** | organic carbon density (kg/m<sup>3</sup>) | IGIP-DIS 2000 |
| **thermcap** | the thermal capacity (only integer 0,1 to 2 J/m<sup>3</sup>/K) | IGIP-DIS 2000 |
| **wiltpont** | the Wilting Point (mm). | IGIP-DIS 2000 |
| **Alt** | elevation (m) | EEA 1990 |
| **bird biodiv** | biodiversity of bird species | Sandel 2011 |
| **mam biodiv** | biodiversity of mammal species | Sandel 2011 |
| **amph biodiv** | biodiversity of amphibian species | Sandel 2011 |
| **bird endnb** | number of bird endemic species | Sandel 2011 |
| **mam endnb** | number of mammal endemic species | Sandel 2011 |
| **amph endnb** | number of amphibian endemic species | Sandel 2011 |
| **connLO** | connectivity of Land Ocean distribution | Gaucherel et al. |
| **shaLO** | Shannon index of Land Ocean distribution | Gaucherel et al. |
| **shaBord** | Shannon index of the fragmentation of continents by | Gaucherel et al. |
| **LCD** | Land Cover evenness index (log transformed) | LADA 2008 (FAO) |
| **HDI** | Human Development Index | UNDP 2012 |
| **UR** | Urbanization Rate | WBO 2012 |
| **pop** | population density (inhabit./km<sup>2</sup>) | GPWFE |

**Figure 1.**
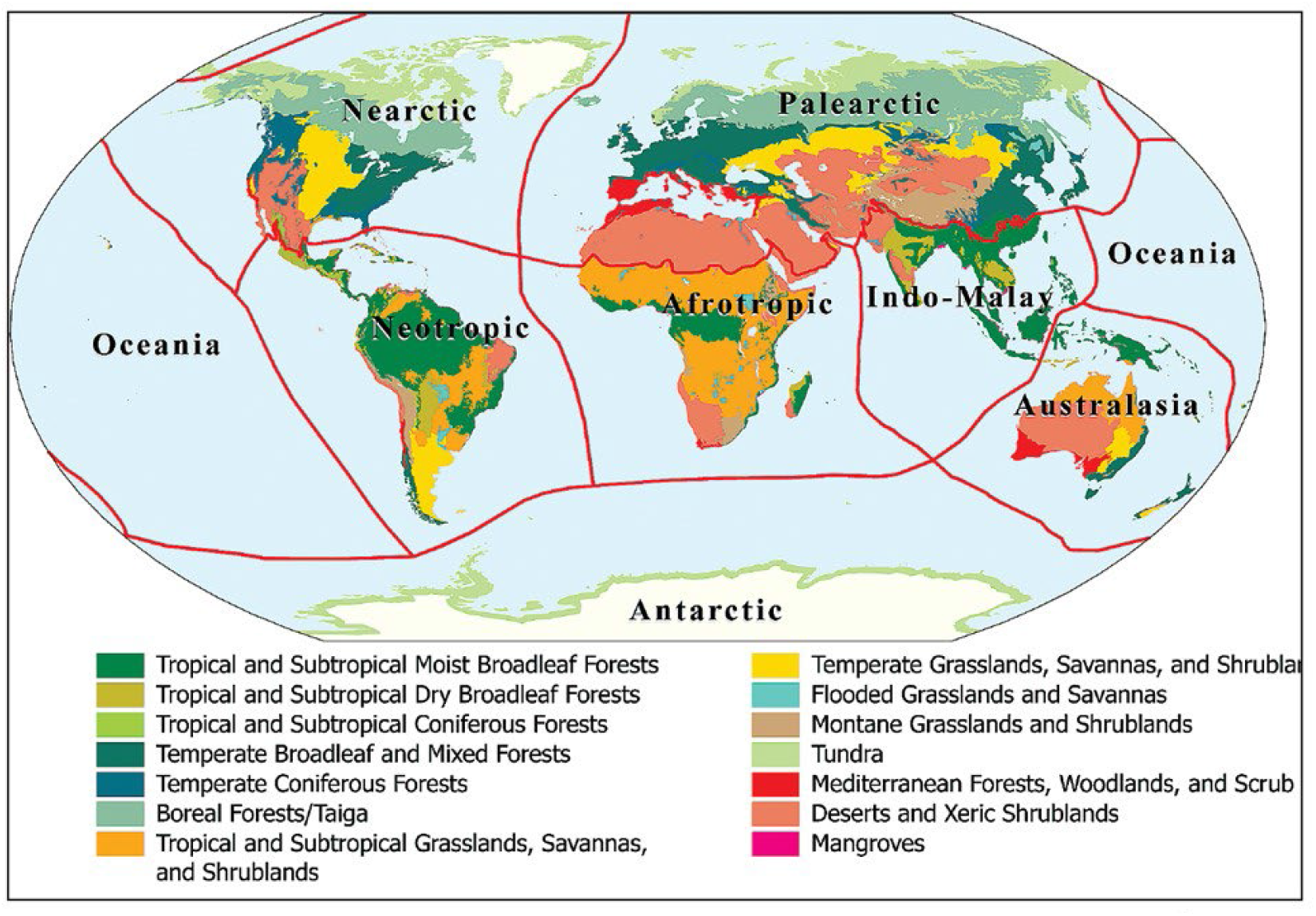
World map of the ecoregions, classified into 14 biomes (Table 1) and eight associated biogeographic realms [16].

It led to 43 variables (climatic, soil-related, and biodiversity/endemism indices) from which we computed a second Boolean matrix by thresholding each variable with its planetary median value. So far, human related variables have been avoided, as we expected comparing potential biomes and not realized biomes with their respective human impacts. For each biome, we set each variable to 1 for values greater or equal to the median and to 0 otherwise. The four geographical variables, surfaces and average latitude and longitude locations of each biome have not been included, to avoid a geographical bias, and to consequently validate the computed phylogeny. Data have then been standardized before computing the pairwise Euclidean distances between Biomes. Finally, 133 traits were coded as 0/1 for 16 biomes, and their weights were defined all equal to unity. The biome ‘Lake’ was set as an outgroup for the 15 remaining ingroup biomes. Phylogenetic relationships among biomes were reconstructed under the maximum likelihood optimization criterion as this method has been shown to be robust against differences in rates of change among phylogenetic units, here the biomes [23]. A F81 model of exchanges among trait states was used. A Gamma distribution with three discrete categories was also used to account for potential variation in the rate of change among traits [24]. The best tree was identified after tree bisection-reconnection branch swapping on a neighbor-joining starting tree under PAUP* [33], version 4.0a. Confidence on the nodes was estimated after hundred bootstrap replicates under the same settings [34].

## Results

A total of 26 equally likely trees were identified by the maximum likelihood search on the 133 “ecological” traits. The consensus topology of these trees is provided on the figure (Fig. 2). Branch lengths were optimized from this consensus under the maximum likelihood criterion, and are proportional to the number of expected trait changes. Bootstrap percentages higher than 50% (a more robust confidence level, see the bootstrap histogram inserted) are shown close to the corresponding nodes, as usually done in phylogeny analyses. In particular, six nodes over 11 nodes appeared quite robust. They mainly highlight transitions among tropical biomes, which seem to present clear-cut trait compositions.

**Figure 2.**
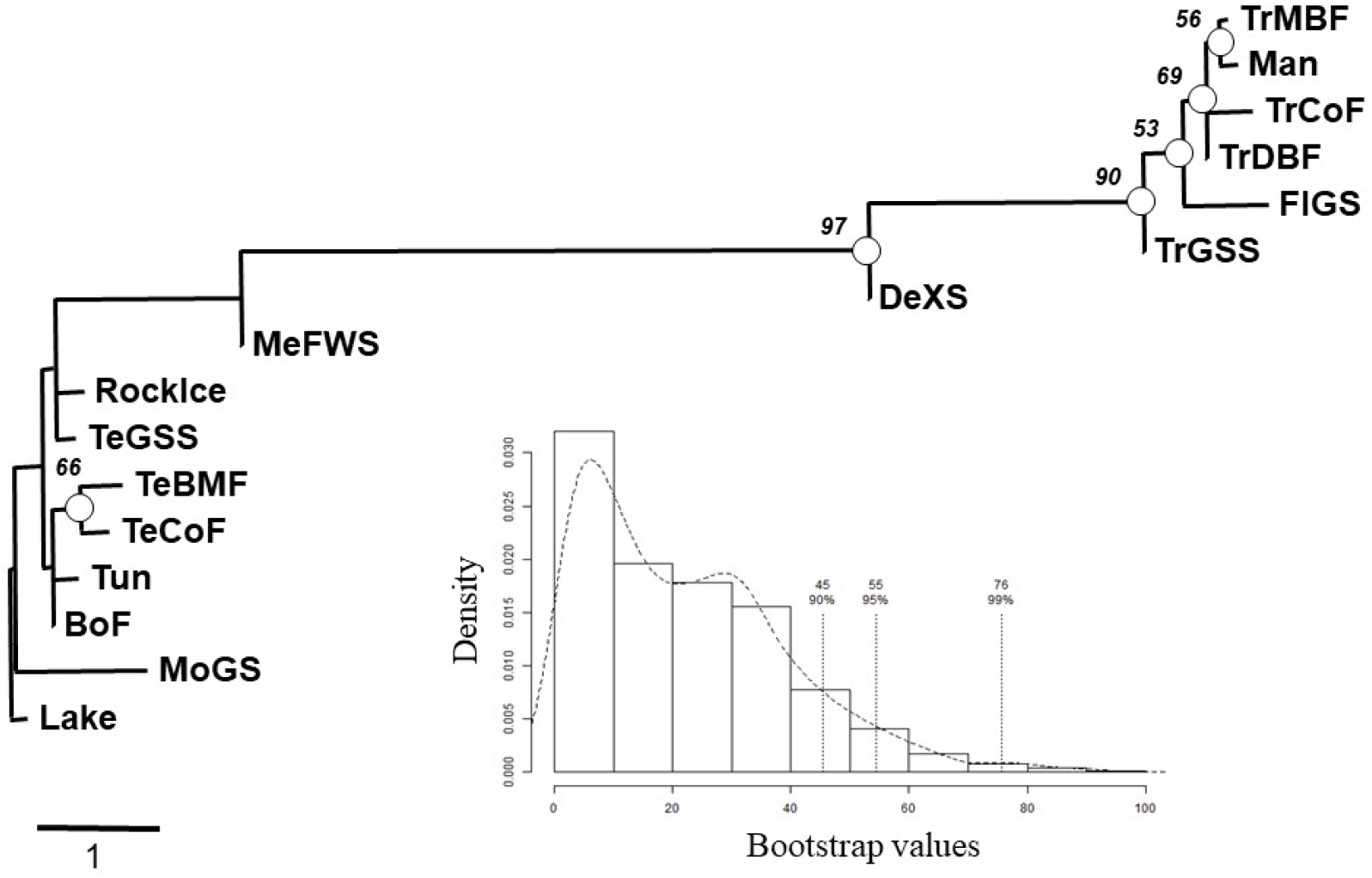
The most probable phylogenetic tree for the 14 biomes showing the consensual topology, with the histogram of random bootstrapping values displayed (insert). The branch lengths are proportional to the number of expected trait changes. Bootstrap percentages higher than 50% are shown close to the corresponding nodes (circles).

Overall, the resulting phylogeny is close to what was hypothesized. In particular, the latitudinal gradient is clearly visible from the node order and their related branches, although geographical variables were not used as direct input variables in the phylogeny computation. The phylogenetic pattern appears robust for tropical biomes as they all relate to bootstrap percentages higher than 50% (Fig. 2, top), and more rarely robust for present-day biomes from higher latitudes (exception for the temperate forest biomes). The others filiations, such as tundra and boreal forests, remain expected although non-significant. Finally, some biomes have a more ambiguous position, such as Mediterranean and xeric ecosystems, as they are located in between the others in this final consensual phylogeny.

## Discussion

The consensual biome phylogeny obtained is not as impressive as we might have expected compared to traditional species phylogenies. This observation comes from the fact that biomes, as for local ecosystems, are understood as composed of biotic, abiotic and anthropic components, in deep interactions [35]. For this reason, the living properties of ecosystems and biomes are still under debate, i.e., whether they might be considered as alive and exhibiting a possible evolution [36]. Yet, the computed biome phylogeny is understandable, which certainly means that bioclimatic variables (and the longitude influence) are driving the observed structure, yet with some departures from climatic traits.

Two reasons may explain why biome lineages could not be modeled linearly (with a vertical transmission): i) biomes are not clear-cut categories as species are supposed to (despite a long-standing debate), and ii) they do not only depend on biological materials and processes building a linear history. Indeed, biomes are made up of abiotic components and processes which regularly blur the phylogenetic signal from the biotic components. Such biomes do not have a clear and stable DNA-like structure submitted to a kind of selection and that would carry the lineage information relating them to each other. In previous studies, it has been proposed that such structure may be made up of the interaction network forming the skeleton of each ecosystem [37-39]. Yet, such a skeleton would not be as stable and unique as the DNA molecule is, thus not playing the role of the clear-cut information storage it plays in biological units. This reminds the old attempt to build a classification of minerals on the basis or Linnea principles, by Mendes da Costa in 1650 [40], a stillbirth attempt due to the erratic (non-vertical) physicochemical processes at play.

As a consequence, physicochemical processes regularly shake biome functional boundaries (and their differences), as well as generate regular changes backward in the phylogeny. This observation is highly similar to genome introgressions and mimic what biologists call recombinations [e.g., 41]. A similar work has been done on the big cat phylogeny, as the phylogenetic tree was blurred by many introgressions [42]. This justifies why we computed the biome phylogeny including biotic traits, the ones which better correspond to a possible phylogenetic context. Our understanding is that the biome phylogeny is accessible based on the species or community the biomes are composed of, only over the relatively short term, i.e., between relatively similar ecosystems [4, 5]. Yet, it seems to be less reliable on the basis of physical traits over a longer term, although we do observe some similarities between neighboring climates and soils. These additional traits remain relevant, and allow computing a longer and more integrated biome phylogeny (Fig. 2), but we ignore how far in the past. The temporal links between abiotic traits involve different mechanisms [40], distinct speeds than for biotic materials, and a priori higher uncertainties. As a consequence, we are not able to identify the ancestor of all biomes, instead of some (more recent, e.g., tropical or colder biome) ancestors of more confidently related biomes (Fig. 2, confidence levels).

Among the processes responsible for biome transitions, some are today well known, such as those based on state successions [e.g., 5, 43], but their interactions, feedbacks and influences over the long term are not [e.g., 8, 44, 45]. In applying phylogenetic tools here, we assumed that all processes at play, including pedological, climatic, and anthropic processes, change randomly on average over the long term. This is still a hot debate to identify the exact contribution of factors responsible of the biome distributions on earth, and of biome transitions [e.g. 7, 8, 46]. Biomes are supposed here to statistically exhibit a Brownian motion of traits and combine independently, and furthermore, at the same speed. As a perspective, more mechanistic models may be develop and tested too [e.g., 12, 15], over the long term and at planetary scale, to confirm this preliminary and correlative analysis. Among biome traits, geological processes have a much lower rate of change than biologically related processes. However, taking into account different rates for these processes did not result in a more realistic biome phylogeny (Not shown). In a preliminary analysis, we additionally used other aquatic biomes, and another analysis based on expert knowledge has been tested, but we found no significant relationships in these other phylogenies.

When computing the phylogeny, we tested the robustness of the proposed phylogeny by varying the speed rate (weights) at which traits may change, with higher rates for biotic traits than for abiotic ones, ultimately resulting in a similar phylogeny (not shown). The best would be to know at which rate each (category of) trait would vary in time. In particular, we tested biological traits varying more rapidly than climatological traits, themselves varying more rapidly than soil related traits. Such changing rates are depending on selection pressure and additional factors, as it is well-known for genetic patterns [47]. Unfortunately, we do not have this information for biomes and proceeded with the minimal assumption that biome traits have a uniform changing rate. Also, we are conscious that such phylogenetic trees are grasping fission events only, not merging events [42]. Taking into account for more complex dynamics of traits would be necessary and is a next step in building a reliable biome phylogenetic tree.

Tropical biomes are clearly differentiated from other biomes (Fig. 2, confidence levels). They are separated mainly due to biological and soil related traits, thus responsible for changes in biome functioning. In a way, such changes appear similar to those identified in ecosystem changes and already identified as the ecosystem development [36, 48]. This may be explained by the biological traits that indeed present a clear-cut phylogeny. We are not sure why tropical biomes and their phylogeny are better identified, but it may be due to the fact that tropical biomes have been less perturbed and remained ice-free during the last glacial periods [11, 19, 49]. In addition, they remained in average at roughly similar (low) latitudes and maintained longer in average. For all these reasons, we conjecture that their phylogeny appeared easier to identify.

The biome phylogeny reveals a biome common ancestor, thanks to the method used, but we are not claiming that this ancestor has ever existed. Rather, we think that ancestors (plural) of terrestrial biomes probably gathered some of the properties (traits) listed in our ancestor phylogeny. Many other studies have used the power of phylogenetic trees without systematically looking for a common ancestor [21, 22]. However, it is reasonable to describe such ancestors as having a lower species diversity, and aquatic traits. It is believed that terrestrial biomes emerged from coastal waters and gradually developed into what we observe today [50, 51]. With more data available on aquatic (i.e., fresh, brackish and salt waters) properties [e.g., 49], we could have computed a single global phylogenetic tree for both terrestrial and aquatic biomes and scrutinize this ancient transition. As for ancestors, we recall that the proposed biome phylogeny is not claiming that we identified the exact history of terrestrial biomes on Earth, rather than one of the many possible histories of biomes. It would be relevant to model biome transitions at large spatial and temporal scales for exploring the possible pathways of the biomes and Earth history [e.g., 52, 53]. Despite this attempt to adopt an integrated view of biome transitions, the phylogeny is more relevant in providing steps of transitions between closely related biomes in the overall succession.

As in many other phylogenies, the biome phylogeny computed here is a clear approximation of the “dominant” biome successions observed on Earth. Biomes are not exhibiting reproduction as cells or organisms, but they also exhibit a kind of persistence, trait heritability and lateral exchanges. It remains a hot debate today to identify a potential “evolution” for ecosystems, and thus for biomes. The proposed phylogeny here brings a weak yet relevant argument for broad and long-term evolution of biomes and ecosystems.

## Acknowledgments

We thank here Romain Frelat for his help in the early analysis of this paper, plus Emmanuel Paradis and Emmanuel Douzery for some early trials of biome phylogeny computations.

## Policy statements

This work received no funding. Ethics declaration: not applicable. Consent to Publish declaration: not applicable. Consent to Participate declaration: not applicable.

